# BactMentha: a database of annotated bacteria-host protein interactions

**DOI:** 10.64898/2026.09.29.755270

**Authors:** Lou Bergogne, Mégane Boujeant, Marta Iannuccelli, Aurélie Bergon, Jaime Fernandez-Macgregor, Christine Brun, Luana Licata, Andreas Zanzoni

**Affiliations:** Aix Marseille University, INSERM, TAGC, UMR1090, Turing Centre for Living Systems, Marseille, France; Department of Biology, University of Rome Tor Vergata, Rome 00133, Italy; Aix-Marseille University, CNRS, AFMB, UMR 7257, Turing Centre for Living Systems, Marseille, France; CNRS, Marseille, France

## Abstract

Bacterial pathogens ensure their successful dissemination by employing specialized secretion systems to deliver a wide range of virulence proteins that interact with specific host proteins to modulate the response during infection. These interactions are often mediated by host-like elements, like short linear motifs, which can interfere with host interaction interfaces, enabling bacterial secreted proteins to compete and displace native partners of the targeted proteins. Several resources provide access to bacteria-host protein-protein interaction data. However, some are neither fully functional or up to date. Moreover, most of these databases lack virulence protein annotations, thus hampering hypothesis formulation based on protein interaction analyses. In this context, we have developed BactMentha, a unified resource that integrates bacterial-host protein-protein interactions for three organisms (human, mouse, and rat) with virulence protein annotations and bacterial strain clinical relevance information. BactMentha provides easy access to available experimentally identified binding regions of the interaction partners, complemented with putative interaction interfaces based on predicted structure complexes and on the detection of host-like elements on bacterial protein sequences interacting with human proteins. BactMentha currently contains 11 776 unique protein interactions between 2970 bacterial proteins, of which 684 have at least one virulence annotation, and 5088 host proteins. Experimentally identified binding regions are available for 532 bacteria-host interactions, most of which involve human proteins (440 interactions). Predicted interaction interfaces enabled a 3.7-fold increase in known and putative binding regions in the bacteria-human protein interaction data. By providing easy access to functionally annotated and clinically relevant protein interaction data, enriched with experimentally and predicted interaction interfaces, BactMentha can be instrumental in formulating novel hypotheses and speeding up wet lab research on the molecular mechanisms underlying pathogenic bacterial infections.

## INTRODUCTION

Bacterial pathogens interact with their hosts to reach an advantageous niche and ensure their successful dissemination by interfering with important host-cell processes like membrane and cytoskeleton dynamics, immune response, or cell death/survival (1–3). To achieve this, they employ specialized secretion systems (4) to secrete and deliver a wide range of virulence proteins (e.g., adhesins, effectors) (5), which interact with specific host proteins to modulate the response during infection. For instance, adhesins bind host-cell membrane receptors to facilitate colonization and invasion (6), whereas effectors translocate directly into the host cell *via* ‘needle and syringe’-like secretory systems, such as type-3 and type-4 secretion systems (T3SS and T4SS), to directly manipulate intracellular pathways (3). These interactions are often mediated by host-like elements that resemble host components in form and function, such as targeting sequences, enzymatic activities, and protein-protein interaction modules, *i*.*e*., globular domains and short linear motifs (SLiMs) (7,8). These elements can interfere with host interaction interfaces, enabling bacterial secreted proteins to compete and displace native partners of the targeted proteins, thus representing an effective interaction perturbation strategy as recently shown in the case of the type-3 *Chlamydia* effector Tri1 that interacts with the human TRAF7 protein (9).

Over the years, interaction discovery experiments permitted the identification of hundreds of bacteria-host protein-protein interactions (PPIs). This data is scattered in different databases, such as IntAct (10) and BioGrid (11), and can be readily gathered via programmatic access (12). IntAct serves as the central IMEx repository, consolidating interaction data from member databases including MINT (13) and UniProt (14). To more easily navigate pathogen-host PPIs, few specialized databases have been developed to easily provide pathogen-host PPIs, including HPIDB (15), PHISTO (16), and, more recently, InTxDB (17). However, these resources are neither fully functional (HPIDB) nor updated (PHISTO), nor available for download (HPIDB and InTxDB). Notably, all data previously available in HPIDB is now accessible via IntAct. In addition, most of these databases lack virulence protein annotations, thus hampering hypothesis formulation based on protein interaction analyses. InTxDB is an exception, as it contains interaction data between bacterial secreted effectors and human proteins. To overcome the current limitations of these resources, we developed BactMentha, a unified resource that integrates, through a fully automated workflow, bacteria-host PPIs for three organisms (human, mouse and rat) from the IMEx consortium databases (18) with annotations from BastionHub (19), the Virulence Factor Data Base (VFDB) (20), cellular localization annotations from UniProt (14), and clinically relevant bacterial strain information from the World Health Organisation (WHO) and the UK Health and Safety Executive (HSE), through the Pathogens Portal (21). Importantly, BactMentha provides easy access to available experimentally identified binding regions of the interaction partners, which are reported as biological features in IMEx-annotated interaction data. These experimental annotations are complemented with putative interaction interfaces predicted using both sequence-based and structure-based tools: the *mimic*INT workflow (22), which detects host-like elements in bacterial proteins, and AlphaFold3 (23), respectively. Overall, by providing a bacteria-host PPI dataset enriched in virulence protein and interaction interface annotations, BactMentha is a valuable resource to study bacteria-host relationships. In particular, it enables the investigation of the molecular mechanisms potentially involved in host-cell network subversion by pathogenic bacteria.

## MATERIAL AND METHODS

### Literature curation

We performed a targeted curation effort aiming at curating pathogen bacteria-host protein-protein interactions, focusing on missing interactions such as those involving pathogen secreted effectors. MINT expert curators extracted relevant information from peer-reviewed articles and annotated them following the detailed IMEx curation model (24) and using a standardized controlled vocabulary (25). This systematic curation work has produced over 3700 new binary interactions extracted from almost 70 articles.

### The BacMentha workflow

Protein interaction data gathering and annotation rely on a computational pipeline, the BactMentha workflow (**Figure 1**), which is written in Python (https://www.python.org/) and uses the Snakemake workflow manager for automated execution (26). It consists of three main steps: *(i)* the bacteria-host protein interaction data collection; *(ii)* the functional and clinical/safety annotation of bacterial proteins and strains, respectively; and *(iii)* the protein interaction interface prediction. In the first step, the BactMentha workflow retrieves the available PPIs for three host organisms (human, mouse, and rat) from the International Molecular Exchange (IMEx) Consortium databases (24) via the PSICQUIC web service (12). Only PPIs associated with a valid PubMed identifier are kept. Next, for each host organism, the PPI data is filtered to retain only the interactions with bacterial proteins by checking the taxonomy identifier of the host interacting protein. The resulting dataset is saved in a MITAB 2.7 formatted file (12), which is further parsed to extract relevant fields, including, when available, experimentally identified binding regions and mutation data impacting protein interactions. The workflow also retrieves the descriptions and the sequences of all bacterial and host proteins in FASTA format via the UniProt API (27). The second step consists of three annotation tasks. In the first, the workflow assigns functional annotations to the bacterial proteins in the BactMentha dataset by performing BLAST (28) pairwise sequence comparisons against reference sequences automatically taken from BastionHub (19), a database of translocated effectors, and the Virulence Factor Database (VFDB) (20). The functional association is inferred only if the two proteins share at least 30% sequence identity over at least 75% of alignment coverage. This step is necessary as we often observed that bacterial proteins present in IMEx consortium databases either do not belong to the reference strains present in BastionHub/VFDB, but rather to closely related strains or isolate, or do not have UniProt identifiers. The chosen thresholds represent a good compromise to confidently annotate these proteins. In the second task, to provide additional biological context to both interacting proteins, available cellular localization annotations and their corresponding Gene Ontology Cellular Component (CC) terms are retrieved via the the UniProt API (27). For membrane-spanning proteins, the topological class or type is also gathered. In the third task, the bacterial strains in the dataset are annotated with available priority categories (i.e., critical, high, and medium) according to the 2024 World Health Organization (WHO) priority pathogens list (29), and with hazard group classification from the 2023 Health and Safety Executive (HSE) Approved List of Biological Agents (https://www.hse.gov.uk/pubns/misc208.pdf). In the third step, the BactMentha workflow integrates the *mimic*INT pipeline (22) to predict putative interaction interfaces by exploiting known domain-domain and SLIM-domain interaction templates (30,31). The prediction procedure is run only on the set of bacteria-human interactions using *mimic*INT default parameters (22) and first checks whether any of the bacterial proteins contain at least one domain or SLIM for which an interaction template is available. Secondly, if the interacting human proteins contain the cognate domain, it infers the interaction interface. All gathered interaction data, functional and clinical annotations, as well as *mimic*INT interface predictions, are stored in tab-delimited text files that are automatically loaded in the BactMentha database.

**Figure 1.**
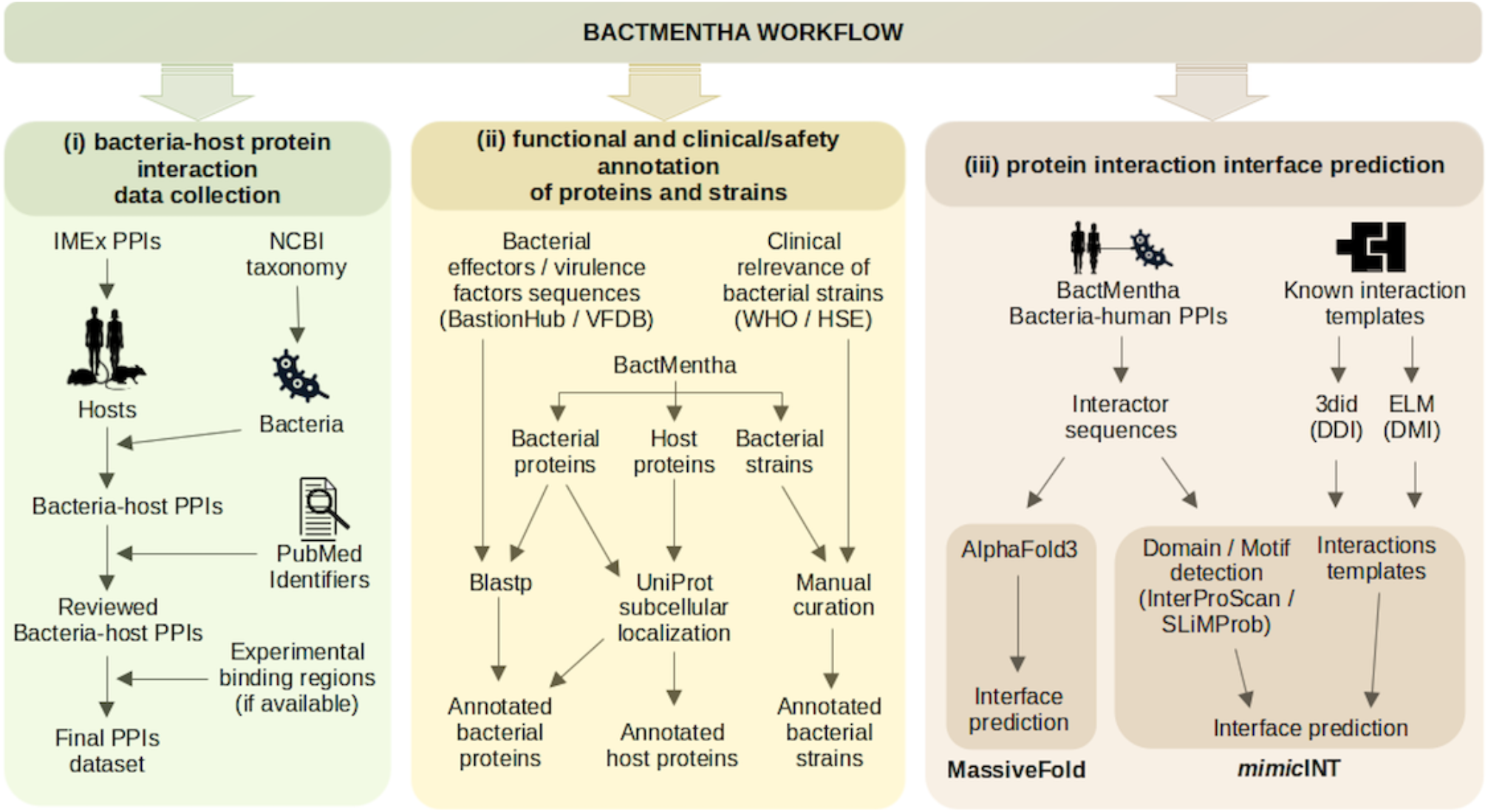
The BactMentha Workflow. (i) Host protein-protein interactions (PPIs) are gathered from the IMEx consortium databases and are filtered using NCBI bacteria taxonomy to keep exclusively bacteria-host PPIs. Only interactions associated with a PubMed identifier are retained. Available experimentally detected binding regions are also retrieved. (ii) BactMentha bacterial protein sequences are functionally annotated by sequence comparison to known bacterial effectors and virulence factor sequences from BastionHub and the Virulence Factor Database. Available subcellular localization annotations are retrieved from UniProt for both bacterial and host proteins. Bacterial strains are annotated based on the priority list from the World Health Organization and hazard groups by the UK Health and Safety Executive. (iii) For the bacteria-human dataset, the domain-domain interaction (DDI) and domain-motif interaction (DMI) interfaces are predicted using the *mimic*INT workflow, and complex structures are predicted by AlphaFold3.

### AlphaFold-based interface prediction

We have prioritized a subset of interactions for AF3 predictions by selecting clinically relevant strains (i.e., hazard group 2 or 3) encompassing different infection strategies (i.e., extracellular/adhesive or intracellular) and secretory system types (e.g., T3 or T4). We used AlphaFold3 (AF3) (23) via the MassiveFold workflow, which is optimized for GPU cluster usage (32), setting the maximum template date to 2024-11-28 and the number of diffusion samples equal to 5. Starting from AF3 output, for each of the five ranked predicted models per interaction, the BactMentha workflow retrieves an computes relevant quality metrics: the average PLDDT value for each interactor chain, the pTM and iPTM score calculated by AF3, the pDockQ2 score (33) and its confidence label (i.e., Low, Acceptable, Medium or High), the number of inter-chain contacts and potential clashes computed following the criteria used in previous works (34,35). As above, this information is stored in tab-delimited text files and automatically loaded in the BactMentha database.

### Database and website implementation

The BactMentha database has been created in a Docker environment for easier deployment and maintenance. We used Docker *(https://www.docker.com/,version27.5.1)* and Docker Compose *(https://docs.docker.com/compose/,version1.29.2)* to enable communication between four different containers. One of the containers provides the open-source object-relational database system PostgreSQL *(https://www.postgresql.org/)* and is created from the official Docker image (*version 13) available on* DockerHub *(https://hub.docker.com/)*. A second container is based on a Python image (version 3.9), which provides the communication between the BactMentha workflow and the database container for content update. A third container carries the open-source NginX (version 1.25.4) webserver and reverse proxy software *(https://nginx.org/)* to host the website, which has been implemented using HyperText Markup Language (HTML), Hypertext Preprocessor (PHP), Cascading Style Sheets (CSS), JavaScript, and Asynchronous JavaScript and XML (AJAX) techniques. A final container provides a CGI process manager and PHP extensions (php:7.4-fpm-alpine3.13) to handle user requests and database queries. The BactMentha database website is freely accessible at: hhttps://bactmentha.tagc.univ-amu.fr/.

## RESULTS

### Database content

The BactMentha database currently includes 11 776 PPIs between 2970 bacterial and 5088 host proteins, most of which (11 004, 93.5% of the total dataset) involve 4554 human proteins (**Table 1)**. Among the 2970 bacterial proteins, 683 (∼23.0%) have at least one functional/virulence annotation assigned by sequence comparison (**Figure 2)**. These proteins participate in 3931 interactions (33.4%). The vast majority of the bacteria-mouse and bacteria-rat PPIs involve annotated proteins, 71.2% (535 interactions) and 95.2% (20 interactions), respectively (**Figure 2)**. Most annotations are related to translocated effector proteins, adherence, motility, immune modulation, and other relevant factors important for pathogenicity.

**Table 1.** BactMentha protein-protein interaction statistics. The table reports the overall and by-host content of the BactMentha database in terms of number of binary PPIs, distinct host and bacterial proteins, different bacterial strains and families interacting with each host species, as well as the number of distinct articles describing the interactions (some publications provide interaction data for more than one host).

|  | **Total** | **Human** | **Mouse** | **Rat** |
| --- | --- | --- | --- | --- |
| **Interactions** | 11776 | 11004 | 751 | 21 |
| **Host proteins** | 5088 | 4554 | 521 | 13 |
| **Bacterial proteins** | 2970 | 2914 | 98 | 12 |
| **Bacterial strains** | 102 | 97 | 27 | 6 |
| **Bacterial families** | 39 | 38 | 16 | 4 |
| **Articles** | 265 | 239 | 34 | 14 |

**Figure 2.**
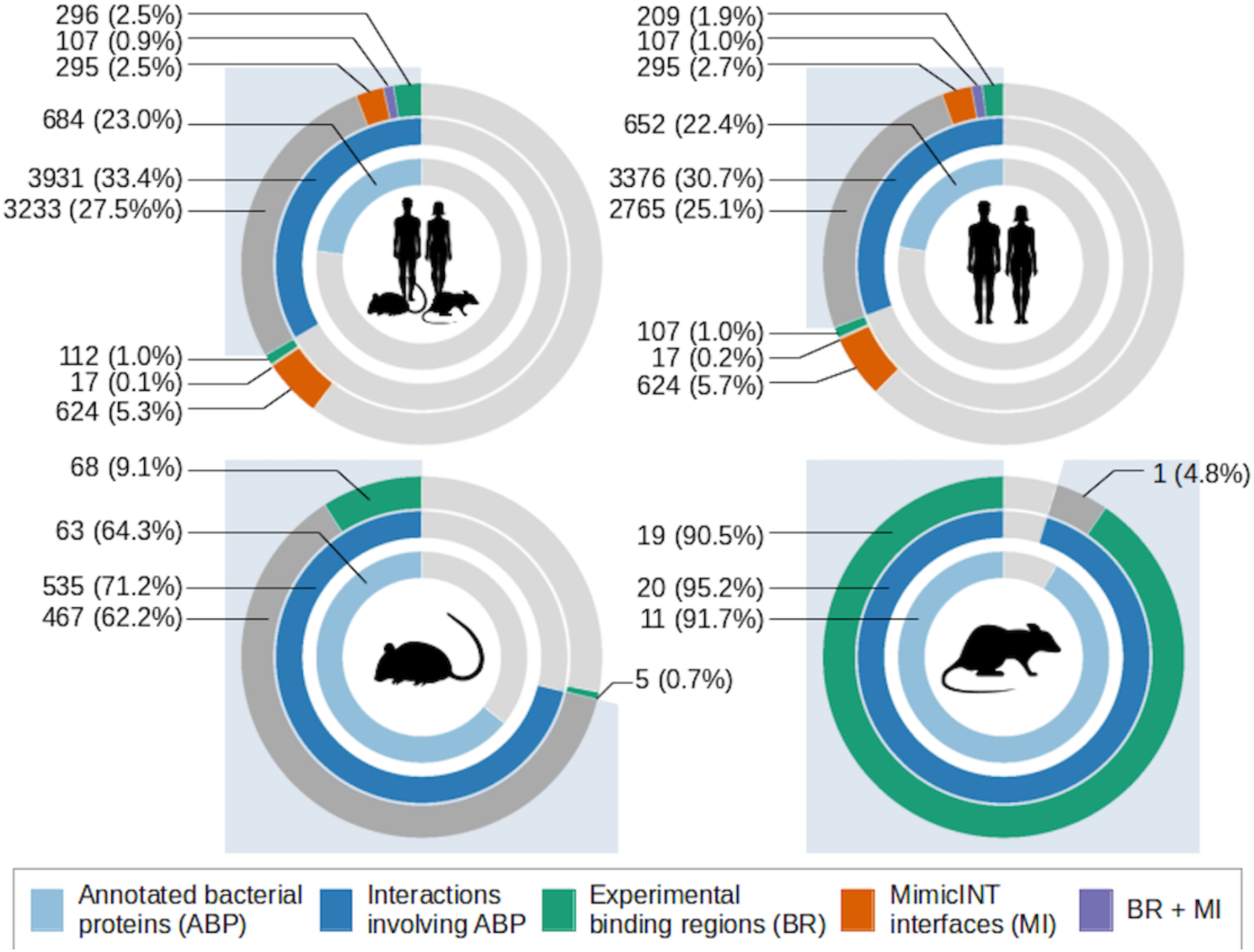
BactMentha annotations of bacterial proteins and interaction binding regions. Coverage of VFDB/BastionHub annotated bacterial proteins (inner circle, light blue), proportion of the interactions in which these proteins participate (middle circle, dark blue + blue background), fraction of interactions with available experimentally identified binding regions only (green), *mimic*INT putative interface only (orange), or both (purple) for the full BactMentha (A), bacteria-human (B), bacteria-mouse (C), and bacteria-rat (D) datasets. Dark grey sectors in the outer circles represent the interactions involving annotated bacterial proteins for which no experimental or predicted binding regions are available. Light grey sectors represent proteins and interactions for which annotations or binding regions are not available.

Only a small fraction of the BactMentha dataset contains information on the experimentally identified binding regions for at least one interaction partner (532 interactions, 4.5%), among which 193 interactions have binding region information for both partners (161 interactions in the human dataset). The integration of *mimic*INT predictions in the bacteria-human PPIs dataset provides a putative interaction interface for 1043 interactions (∼9.4% of the bacteria-human dataset), 124 of which already had experimental binding region information for at least one partner, and 87 of them for both bacterial and human proteins (**Figure 2**). AF3 provides complex predictions for 379 bacteria-human PPIs (3.4%), 61 of which have already an experimentally identified binding regions. The 919 additional interactions for which interfaces have been predicted either by *mimic*INT or AF3, and for which no binding information was available, enabled an increase from 440 to 1641 (∼3.9% to ∼14.9%) the number of bacteria-human PPIs with either experimental or putative binding information. At the scale of the global dataset, this number has increased from 532 to 1733 PPIs, reaching a proportion of ∼14.7% of BactMentha dataset for which molecular binding regions are available.

Among the 102 bacterial strains present in the full dataset, belonging to 39 different bacterial families, 11 are included in the WHO *critical* priority group, including distinct *E. coli* strains, 16 to the *high-priority* group (such as some *Salmonella, Shigella* and *Pseudomonas*), and 7 to the *medium* priority group (like *Haemophilus influenzae* or *Streptococcus pyogenes*). Most of the strains are present in the HSE list of biological agents, namely 86 (84%): 72 strains belong to Hazard Group 2, while 14 are in Hazard Group 3. While both groups include pathogens that can cause human diseases for which a treatment or a vaccine is usually available, Group 3 pathogens can cause severe diseases that may spread to the community.

### Database web interface

The BactMentha database home page provides the user multiple search modalities, such as a quick search query form to interrogate its content via a UniProt identifier, a gene name, an NCBI taxon identifier, or a PubMed ID (**Figure 3)**. The user can also perform complex queries through an advanced search form, which allows combining multiple terms. Alternatively, the user can directly explore the interaction data based on the host organism or access the Browse page. The latter features pre-complied data views based on the WHO priority list and HSE hazard groups and provides direct access to bacteria-human interaction data for which experimental binding regions or predicted interaction interfaces are available. The query results are displayed in a tabular format. The interface allows to dynamically filter the table content with a global free-text search or by column. The user can add or remove columns through the “Show/Hide Columns” button (**Figure 4**). For each interacting pair, the user can display, if available, the bacterial protein (PA) and the cellular components (CC) annotations, the experimentally identified binding regions (BR), and the predicted interaction interfaces by *mimic*INT (MI) and AF3 (AF) by clicking on the corresponding buttons in the last table column (**Figure 4**). The result table can be exported in comma-delimited format, including protein and interaction annotations, or xlsx format with just one click.

**Figure 3.**
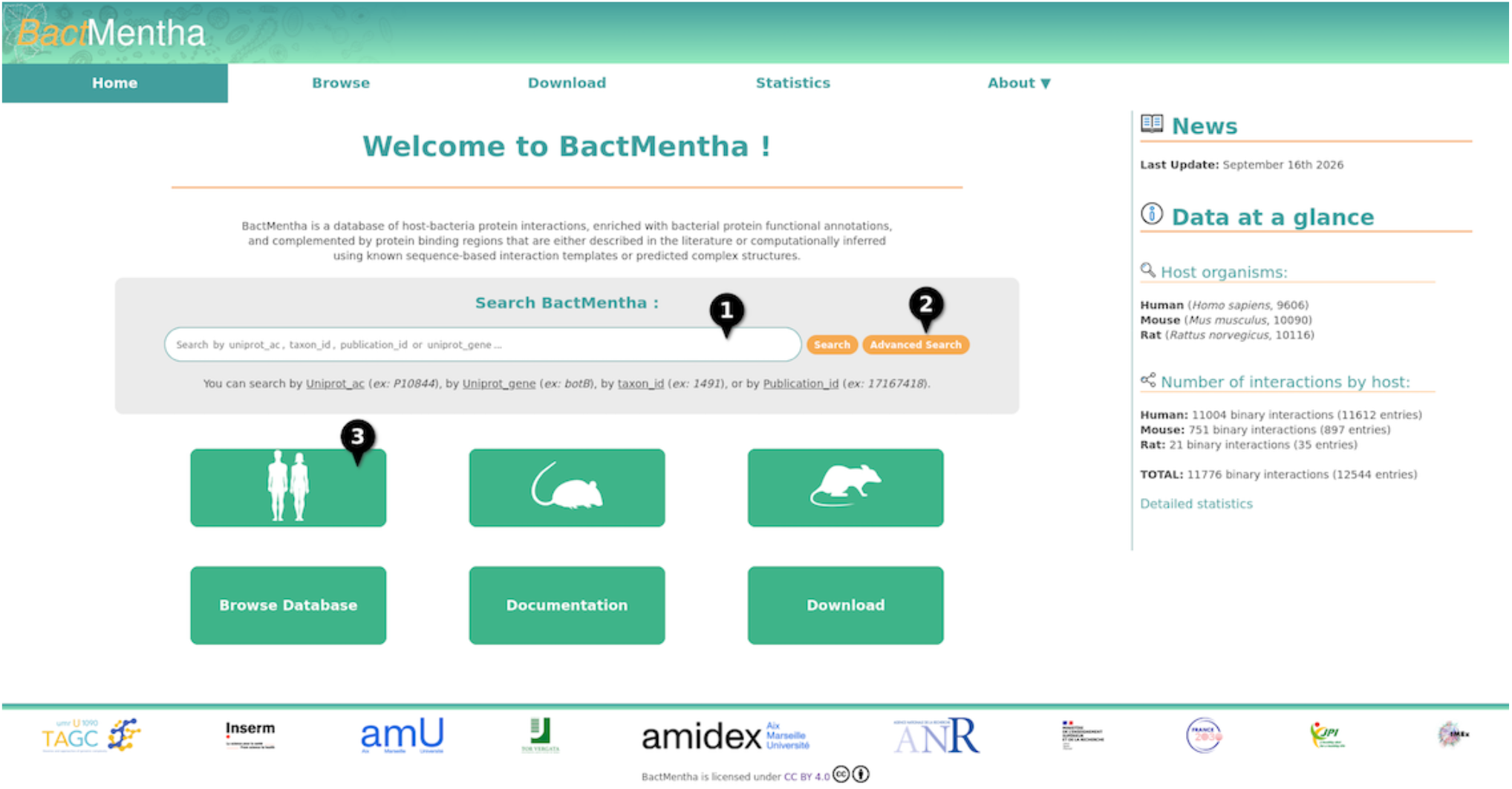
BactMentha website Home page. BactMentha Home page, available at https://bactmentha.tagc.univ-amu.fr/, provides (1) a quick search bar and (2) an advanced search form. (3) The database can be accessed via preselected datasets by host or through several browsing options.

**Figure 4.**
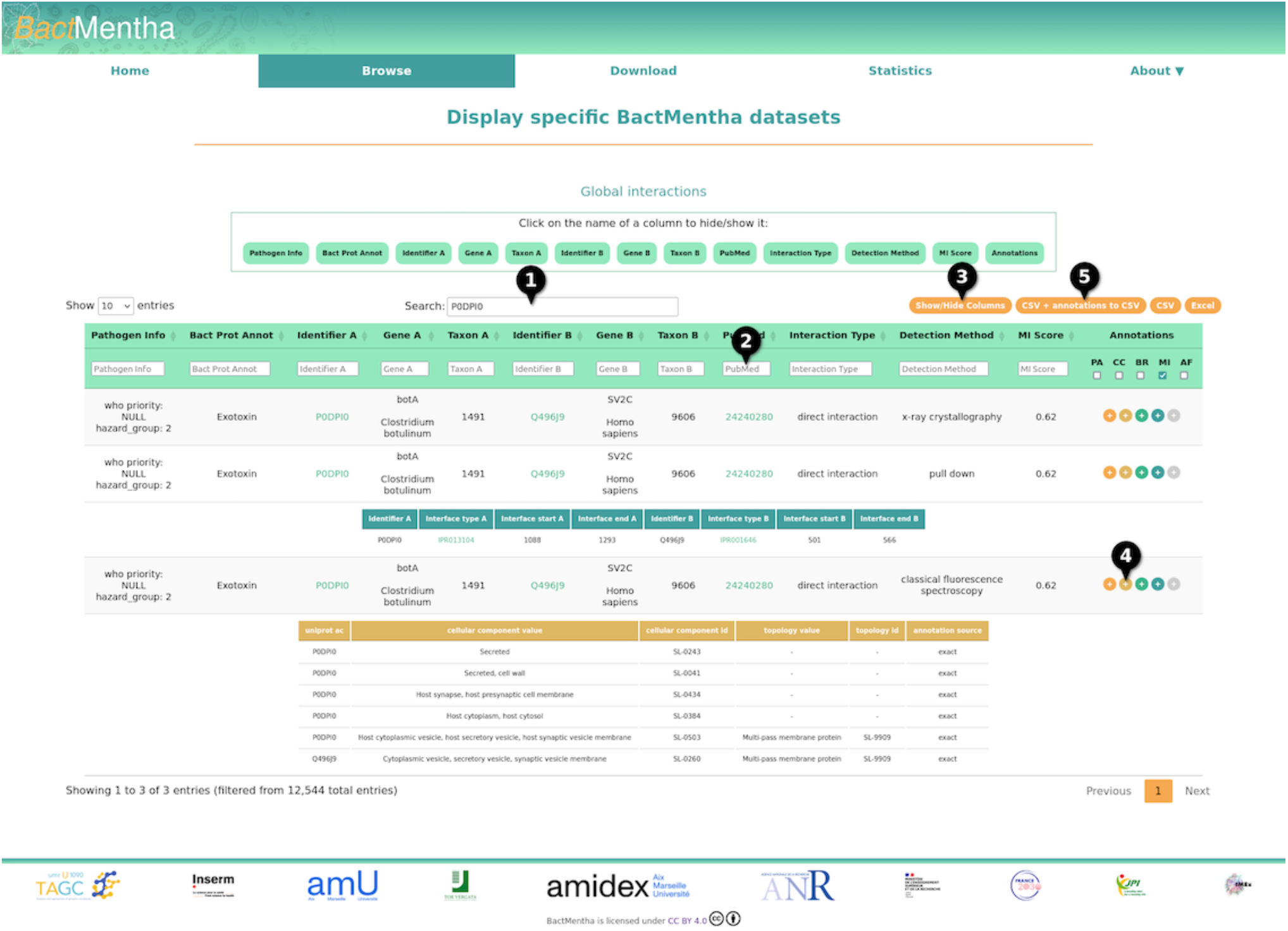
BactMentha result table. Query results are displayed in a table that provides quick filtering options through dedicated (1) whole table search bar, or (2) column-based searches (e.g., organism or interaction detection method) or checkboxes (e.g., protein with annotations or with experimental/predicted binding regions). The user can also select or hide table columns to display by clicking on the (3) “Show/Hide Columns” button. For each interaction, when available, additional information can be displayed by clicking on (4) the annotation column buttons for visualizing the (PA) the bacterial protein annotations, subcellular localization annotations (CC), (BR) the experimentally detected binding regions, and (MI) and (AF) for *mimic*INT and AF3 predicted interaction interfaces, respectively. The results table, along with protein and interaction annotations can be (5) downloaded in comma-delimited or xlsx format.

The user can access *Statistics* and *Download* pages via the navigation bar. On the latter, current and archived BactMentha datasets are available to the community. They include interaction data as well as bacterial protein and interaction annotations in tabular format (*database table* files) together with all the files produced by each step of the BactMentha workflow (*reference data* files). The user can download the full dataset or host-specific data. The Statistics page provides data summaries with a particular focus on protein and interaction annotations.

Through the *About* drop-down menu, the user can access the *Documentation* page, which provides details on data gathering and a user guide, a frequently asked questions section, and the contact page.

### Case studies

*Coxiella burnetii* (strain Nine Mile phase I, HSE Hazard Group 3), an obligate intracellular pathogen causing the Q-fever disease (36), is among the clinically relevant bacterial strains with the most available interaction data. Indeed, BactMentha contains 238 interactions between 20 *Coxiella burnetii* proteins, of which 19 are functionally annotated as effectors, and 221 human proteins. Only one experimentally identified binding region is available for the effector protein CvpF, which interacts with the human small GTPase RAB26 (37). Notably, BactMentha provides at least one *mimic*INT predicted interface for 23 of these interactions between 9 Coxiella *burnetii* and 22 human proteins. For instance, the effector protein CBU0881 (UniProt identifier: Q83D60) interacts with the human DNA topoisomerase 2-binding protein 1 (TOPBP1, UniProt identifier: Q92547), a scaffold protein involved in DNA replication and repair. TOPBP1 also acts as an apoptosis inhibitor by interacting with E2F1, through a BRCT domain, thereby repressing E2F1 transcriptional activity (38). BactMentha interface predictions indicate that this interaction can be mediated by a BRCT-binding motif located on the N-terminus of CBU0881 protein (ELM identifier: LIG_BRCT_BRCA1_1), which can be recognized by TOPBP1 BRCT domains. This suggests that CBU0881, by interacting with TOPBP1, may participate in host cell apoptosis inhibition, as it has been shown for other *Coxiella* effectors (39,40). Another interesting example is the interaction between the effector CBU0937 (UniProt identifier: Q83D09) and the Na(+)/H(+) exchange regulatory cofactor protein (UniProt identifier: Q5T2W1), known also as NHERF3, and coded by the PDZK1 gene. BactMentha provides predicted interfaces between a C-terminal PDZ-binding motif (PBM) detected on CBU0937 and the NHERF3 PDZ domains. Many known bacterial effectors bear functionally relevant PBM in their C-terminal sequences (41,42). NHERF3 is a scaffold protein involved in the regulation of membrane-related processes and vesicle trafficking (43), which are important for the formation and maintenance of Coxiella-containing vacuoles (44). Finally, there is only interaction for which a AF3 predicted complex can be considered of sufficient quality (ipTM = 0.64): the effector protein CBU0175 (UniProt identifier: Q820B6), which contains a serine/threonine protein kinase domain, and the human RNA-binding protein with multiple splicing (RBPMS, UniProt identifier: Q93062). The model predicts that CBU0175 targets the RBPMS RNA-binding domain, which is supported also by *mimic*INT inference. Indeed, RBPMS is a regulator of alternative splicing in vascular smooth muscle cells (VSMCs). *Coxiella burnetii* mainly targets host macrophages (45), but it can also infect other cell types including VSMCs (46). Notably, chronic Q-fever cases mainly present endocarditis or vascular infections (47). This suggests that CBU0175 may interfere with RBPMS splicing regulation in infected VSMCs.

## DISCUSSION AND CONCLUSION

BactMentha provides a unified resource for investigating the potential molecular mechanisms involved in host-cell subversion by pathogenic bacteria. In contrast to other databases, BactMentha integrates bacteria-host PPIs with bacterial protein annotations related to virulence (e.g., secreted effectors, invasins, adhesins) as well as priority and risk classifications for bacterial strains to provide easy access to clinically relevant protein interaction data. In addition, it grants easy access to available experimentally identified binding regions of the interaction partners, which are complemented with putative interaction interfaces predicted by *mimic*INT that detects host-like elements in bacterial proteins. It has been shown that these elements can mimic the host protein interaction interfaces, enabling effectors to compete and displace native partners of the targeted proteins, thus representing an effective interaction perturbation strategy (7,8). Notably, *mimic*INT can identify bona fide interaction interfaces (22,48), thus supporting the relevance of integrating protein interaction data with interface predictions as in the above-described case studies. BactMentha also includes AF3 complex structure predictions, including relevant confidence metrics, for a subset of interactions between clinically relevant bacteria annotated proteins and their human interactors, thus providing structural insights that are complementary to *mimic*INT inferred interfaces (48).

Notably, *mimic*INT predictions can enrich IMEx-curated experimentally binding regions since they very often lack information on the identity of the molecular elements (i.e., domains or short linear motifs) involved in the described interactions. For instance, the TcpB virulence protein from *Brucella melitensis*, a major zoonotic agent causing human brucellosis (HSE hazard group 3), interacts with the human adapter protein TIRAP (49). Structural studies have shown that both TcpB and TIRAP can form dimers through their respective TIR domains (49,50). However, the information is currently absent from the source IMEx databases. Importantly, BactMentha provides the known interface between the two TIR domains, thanks to the *mimic*INT prediction.

The current version of BactMentha focuses on three mammalian hosts (*i*.*e*., human, mouse, and rat) and provides predicted interfaces for one of them (*i*.*e*., human). In the next versions, we will include additional hosts for which protein interaction data with bacterial proteins is available in IMEx databases, such as *Arabidopsis thaliana* (51). Regarding the bacterial protein annotation step, we plan to implement a more sensitive approach based on profile-based sequence searches (e.g., HMM profiles) to improve the functional annotation inference. Moreover, we will extend the *mimic*INT interface predictions to additional bacteria-host pairs for which the presence of interaction elements (*e*.*g*., short linear motifs) has been determined and validated (31). To further increase the coverage of interface-resolved protein interactions, we will also expand the structure-based interface predictions using AlphaFold pipelines to the whole BactMentha dataset with additional confidence metrics (35,52). We will also continue our effort in targeted curation of the scientific literature describing bacteria-host protein interactions to be included in BactMentha.

We believe that BactMentha is an essential resource for researchers studying bacteria-host relationships. By providing easy access to functionally annotated and clinically relevant protein interaction data, enriched with experimentally and predicted interaction interfaces, BactMentha can be instrumental in formulating novel hypotheses and speeding up wet lab research on the molecular mechanisms underlying pathogenic bacterial infections.

## AUTHOR CONTRIBUTIONS

Lou Bergogne: Data Curation, Formal Analysis, Investigation, Methodology, Software, Visualization, Writing – Original Draft Preparation; Mégane Boujeant: Data Curation, Investigation, Methodology, Software. Marta Iannuccelli: Data Curation, Investigation; Aurélie Bergon: Software. Jaime Fernandez-Macgregor: Formal analysis; Christine Brun: Funding Acquisition, Supervision, Writing – Review & Editing. Luana Licata: Data Curation, Investigation, Supervision, Writing – Review & Editing; Andreas Zanzoni: Conceptualization, Data Curation, Formal Analysis, Funding Acquisition, Investigation, Methodology, Visualization, Project Administration, Supervision, Writing – Original Draft Preparation, Writing – Review & Editing.

## ACKNOWLEDGEMENTS

We would like to thank Gianni Cesareni and Jérémie Perrin for their advice and fruitful discussion. We thank Luke Lambourne (Dana-Farber Cancer Institute, Boston, MA, USA) for sharing the code to compute inter-chain residue contacts and potential clashes. This work was granted access to the HPC and storage resources of GENCI at IDRIS under grant 2026-AD010315212R2, using the CSL and A100 partitions of the Jean Zay supercomputer.

